# Filling knowledge gaps on vulnerable electric ray *Narcine timlei* (Elasmobranchii: Torpediniformes: Narcinidae) through ecological and biological studies

**DOI:** 10.1101/2025.09.24.678286

**Authors:** Amit Kumar, Shrutika Raut, Silpa M.S., S. Prakash

**Author notes:** Department of Biological Sciences and Biotechnology, Institute of Chemical Technology, Mumbai (Shrutika Raut).

## Abstract

*Narcine timlei* (Bloch & Schneider, 1801), commonly known as the brown numbfish, recently moved from Data Deficient to Vulnerable in IUCN Red List. However, the biological information on the species is still lacking. In the present study, we provide an integrative assessment of populational dynamics on *N. timlei*. Morphological and molecular analyses ascertain the species identity as *N. timlei*. A total of 331 individuals were examined, comprising 156 males and 175 females, including 13 gravid, bringing an overall sex ratio of 1:1.21. We observed the highest specimens during the post-monsoon season (33.5%), and minimum during summer (18%). Length-weight relationships indicated negative allometric growth (b=2.78). Most specimens fall under the 17-20 cm size class, and a few exceeded 23 cm. Pregnant females carried 2 to 9 embryos. A total of 69 embryos from 13 females were obtained. Embryo length and weight were 5.18 ± 0.22 cm and 2.02 ± 0.36 g, respectively. No correlation was observed between maternal length and fecundity (R^2^ = 0.0036). Male to female embryo ratio was skewed to 1:1.3. The average length of electric organs in adults was 4.30 ± 0.47 cm. Mean electrocyte count for adults was 117.55 ± 14.48, while in embryos it was 123.36 ± 15.9, indicating early developmental establishment of electrocyte structure in *N. timlei*. We strongly recommend long term species-specific monitoring and management measures to ensure conservation of this ecologically significant but often overlooked batoid.

## 1. Introduction

*Narcine timlei* (Bloch & Schneider 1801) (Elasmobranchii: Torpediniformes: Narcinidae), commonly known as brown numbfish, is the smallest member of the genus *Narcine*. They can be identified by a subtrapezoidal disc, a uniform purplish-brown, brown, or tan dorsal colouration, and wide and broadly rounded tooth bands (De Carvalho, Compagno, and Last 1999; Last et al., 2016). *Narcine timlei* was originally described from the Coromandel Coast, Tamil Nadu, south-east India (De Carvalho, Compagno, and Last 1999). Though the etymology of the species is not explained, it possibly refers to Tamil Nadu, India, where the type locality (Tranquebar) is situated (“Habitat ad Tranquebariam Timlei Malais dicta”) (https://etyfish.org/).

They thrive in the inshore continental shelf at a depth of 5-50 m over soft bottoms, and are distributed widely in the Indo-West Pacific: Arabian Sea (Pakistan) to South-East Asia (India, Thailand, Malaysia, Singapore, Indonesia, Vietnam, and southern China) (Last et al., 2016). Not much biological information exists for this species, except that they can reach a total length of 36 cm, males mature at 14 cm, and birth size is 6 cm. They possess electric organs which generate electric discharges to stun prey and to defend themselves. Based on the estimated model predictions, it was suggested that they prefer a temperature range of 21.9-28.6 □. They belong to trophic level 3.1 ± 0.3 (https://www.fishbase.se/summary/Narcine-timlei). Using the information of closely related species, *Narcine entemedor*, the generation length of *N. timlei* has been inferred as 4 years (VanderWright et al., 2021).

*Narcine timlei* is a slow-growing species with prolonged gestation periods and low fecundity. These life-history characteristics make them susceptible to overexploitation (Bhagyalekshmi and Kumar 2021). Across the range of its distribution, *Narcine timlei* is subjected to high fishing pressure due to unsustainable fishing practices like trawlers (VanderWright et al., 2021). Even though this species is not consumed by humans and fishers are not conducting target fishing, they are commonly caught bycatch batoids in the industrial and artisanal fishing (Raut, Prakash, and Kumar 2023; Bhagyalekshmi and Kumar 2021; VanderWright et al., 2021). The brown numbfish is mostly discarded due to its small size or processed as a part of the unsorted bycatch for fish meal production (Raut, Prakash, and Kumar 2023; VanderWright et al., 2021).

The International Union for Conservation of Nature (IUCN) status for *N. timlei* was recently revised from “Data Deficient” to “Vulnerable” (VanderWright et al., 2021). However, the change in IUCN status has not shown any changes in the conservation situation on the ground. They are still a very common bycatch in the mechanised and artisanal fisheries, at least in the southeast coast of India (Bhagyalekshmi and Kumar 2021; Kumar and Prakash 2023). IUCN assessment report 2021 highlighted the lack of species-specific knowledge for *N. timlei* (VanderWright et al., 2021). The assessment report was based on the decreased landings of overall elasmobranchs across the range of *N. timlei*, and it was suspected that a population reduction of almost 50% over the past three generation lengths (12 years) due to exploitation, leading to the vulnerable status of the species. Therefore, it is imperative to gather species-specific data of brown numbfish for their effective conservation. Hence, in this study, we assessed the population structure and reproductive biology of *N. timlei*.

## 2. Materials and methods

### 2.1 Study Location and Sample Collection

Covelong is an artisanal fishing village located 40 km south of Chennai, Tamil Nadu, India. Covelong fishermen communities use gill net, bottom-gill net techniques, and sometimes use hook & line, shore seine and other fishing methods. Fishing activities happen every day by deploying the nets in the afternoon and evening and withdrawing them the next day early morning. They usually fish at a depth of 0-20 m, up to 5 km from the shoreline (A. Kumar, Vinuganesh, and Prakash 2021).

We collected and analysed Individuals of *N. timlei* from the bycatch at the Covelong fish landing centre (12°47’31’’N; 80°15’04’’E) during 2021-2023 in different seasons: premonsoon (July-September), monsoon (October-December), post-monsoon (January-March), and summer (April-June). The specimens were photographed in the field using a Canon PowerShot G16 digital camera and weighed using a digital weighing balance. Whenever possible, we collected the specimens and brought them to the Centre for Climate Change Research facility at Sathyabama Institute of Science and Technology, Chennai, for further detailed studies.

### 2.2 Species Identification Through Integrative Taxonomy

The collected specimens were identified by analysing morphological and meristic characters following published literature (De Carvalho, Compagno, and Last 1999; Ahmad et al., 2013). A tissue sample from a representative specimen was collected and stored in 95% ethanol for molecular identification. Total genomic DNA was extracted using OMEGA BIO-TEK E.Z.N.A. Blood & Tissue DNA Kit, USA, following the manufacturer’s protocol. PCR amplification of mitochondrial cytochrome oxidase subunit I (COX1) gene was conducted using FishF1 & FishR1 primers (Ward et al., 2005) following the PCR reaction mixture and thermal profile published elsewhere (A. K. Kumar and Prakash Prakash 2023). Purified PCR product was sequenced on ABI Prism 3730 Genetic Analyser based on BigDye Terminator Chemistry. Sequence chromatograms were visualised, edited, and contigs were prepared in BioEdit (Hall and Others 1999). The sequences were then compared to published representative sequences of the genus *Narcine* as the ingroup and *Narke dipterygia* as the outgroup terminal. The phylogenetic tree was inferred through the Maximum Likelihood method with the General Time Reversible substitution model on the IQTREE Web Server (http://iqtree.cibiv.univie.ac.at/) (Trifinopoulos et al., 2016).

### 2.3 Population Dynamics

To understand the population structure, brown numbfish were collected seasonally. We identified the sex of the specimen based on the presence of claspers on the pelvic fins of the males, and the sex ratio was calculated in the different seasons. We analysed the Length-Weight relationship. Briefly, length and weight were measured using a vernier calliper and a digital weighing balance, respectively. Length-weight relationship in the form of W= aL^b^ was obtained through linear regression of log_10_ (W) versus log_10_ (L) (Froese 2006). The data analyses were conducted in Microsoft Excel 2010. Disc width was measured as a proxy of body size for batoids.

Further, gravid females were collected for studying embryos. Embryos were also analysed for sex determination, and their length-weight relationship was also observed. The correlation between the total length of the mother and the number of embryos was analysed by employing a regression test.

### 2.4 Electric Organ Morphometrics

The length and width, and number of electrocyte cells in each electric organ were determined under a stereo zoom microscope (Nikon SMZ25, Japan) equipped with a digital camera, and measurements were taken using software (NIS-Elements V5.30, Nikon, Japan).

## 3. Results and Discussion

### 3.1 Morphological and Molecular Identification

The morphological and meristic characters analysed on the collected specimens fall within the ranges reported in earlier taxonomic works (Carvalho, Compagno, and Last 1999), Ahmad et al., 2013; Last et al., 2016), thus confirming their identities as *Narcine timlei*. Key characters of the representative individuals of *Narcine timlei* are shown in Table 1 and Figure 1.

**Table 1.** Proportional measurements of *N. timlei* (n=4) expressed as % of standard length.

|  | *Narcine timlei* |
| --- | --- |
| **Standard length** | 14.63 ± 1.32 |
| **Disc Width** | 62.98 ± 5.21 |
| **Disc length** | 54.89 ± 1.9 |
| **Inter orbital** | 8.4 ± 0.69 |
| **Length to 1<sup>st</sup> dorsal fin** | 74.87 ± 2.27 |
| **Length to 2<sup>nd</sup> dorsal fin** | 86.9 ± 2.08 |
| **Tail base** | 14.77 ± 1.26 |
| **1<sup>st</sup> dorsal fin base** | 7.58 ± 0.91 |
| **2<sup>nd</sup> dorsal fin base** | 7.67 ± 1.01 |
| **Tail width** | 14.77 ± 1.17 |
| **Tail length** | 50.51 ± 4.05 |
| **1<sup>st</sup> dorsal fin height** | 10.37 ± 1.9 |
| **2<sup>nd</sup> dorsal fin height** | 11.41 ± 1.86 |
| **Spiracle width** | 2.5 ± 0.45 |
| **Spiracle length** | 3.88 ± 0.57 |
| **Eye diameter** | 1.9 ± 0.26 |
| **Inter spiracle width** | 7.61 ± 0.15 |
| **pre pelvic** | 41.74 ± 23.85 |
| **pre anal** | 61.17 ± 3.64 |
| **Snout, preorbital** | 17.2 ± 1.55 |
| **Snout, preoral** | 15.05 ± 0.84 |
| **Mouth width** | 11.41 ± 1.89 |
| **Inter nasal** | 7.07 ± 0.87 |
| **Inter 1<sup>st</sup> gill slit** | 16.97 ± 0.89 |
| **Inter 5<sup>th</sup> gill slit** | 9.84 ± 0.4 |
| **Snout to 1<sup>st</sup> gill slit** | 26.9 ± 1.69 |
| **Snout to 5<sup>th</sup> gill slit** | 38.06 ± 2.97 |
| **Pelvic fin base** | 13.6 ± 2.59 |
| **Pelvic fin height/length** | 20.89 ± 6.3 |
| **Total length** | 121.33 ± 3.01 |
| **Clasper base** | 7.79 ± 0.61 |
| **Clasper length** | 15.74 ± 1.15 |

**Figure 1:**
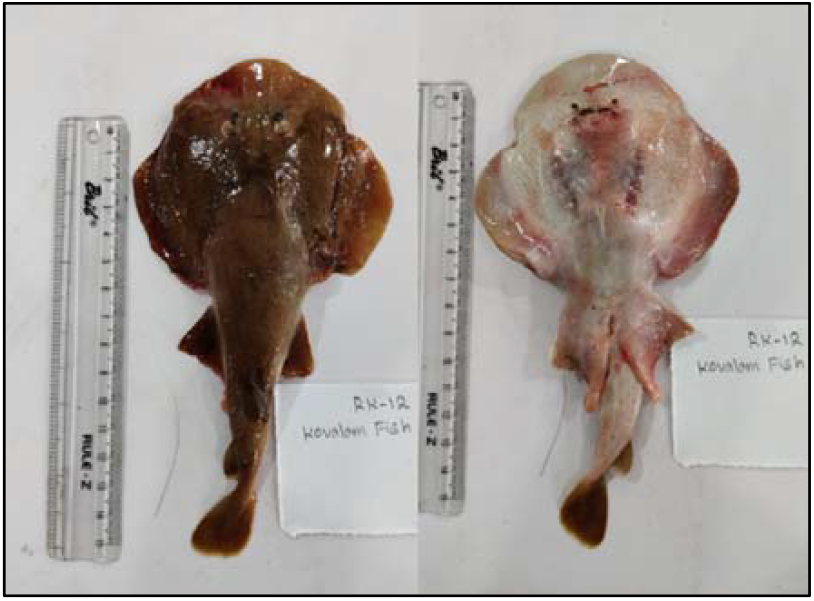
Dorsal and ventral view of the male representative specimen of *Narcine timlei* The species identity was further ascertained through mitochondrial cytochrome oxidase subunit I phylogenetic analysis. The maximum Likelihood tree clustered the samples from the present study with conspecifics from the public database (Figure 2).

**Figure 2:**
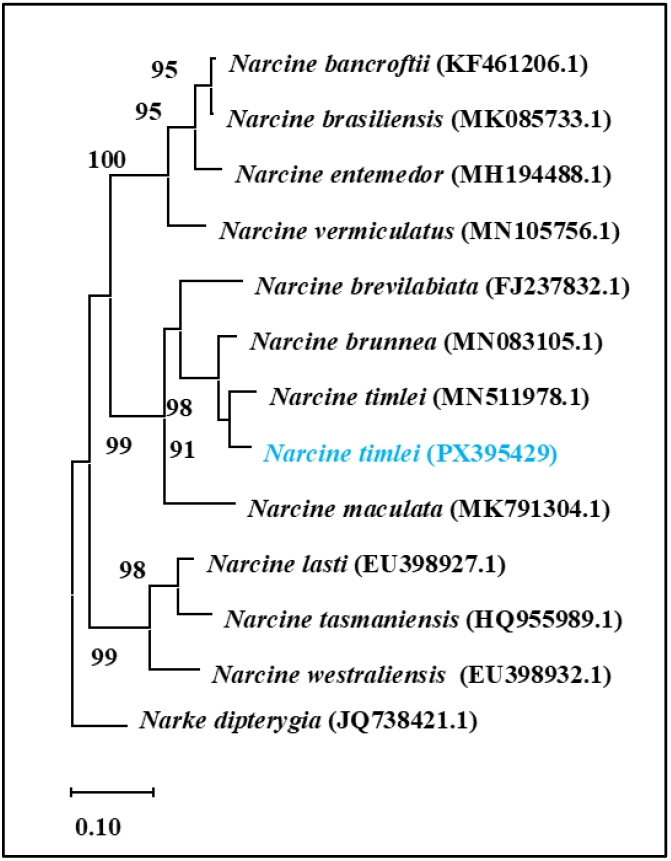
Maximum Likelihood phylogenetic tree of *Narcine timlei* based on mitochondrial gene Cytochrome Oxidase subunit I. The numbers near the branches indicate bootstrap values. GeneBank accession numbers are given in the tree next to the species name. The sequence generated in this study was highlighted in blue.

Earlier studies have shown that morphological identification alone can be confusing, as several characters are similar to congeneric species such as *N. brunnea* (Carvalho and White 2016). Hence, integrative taxonomy combining morphology with molecular barcoding adds robustness to the species identification, therefore providing confidence in the dataset obtained for the species-specific ecological and reproductive analysis.

### 3.2 Population Dynamics

During the present study, a total of 331 individuals of *N. timlei* were analysed for the length-weight relationships, gender, and seasonal availability. Out of 331 individuals, 156 were male and 175 were female, including 13 pregnant women. We found a sex ratio of 1: 1.21 for male: female. While this deviation is not extreme, it suggests an understated shift in population characteristics and their interplay, possibly due to fishing pressure or seasonal distribution. There are no earlier reports available for the sex ratio for *N. timlei* globally. However, studies on other *Narcine* spp. suggested the sex ratio patterns often vary with age, size and geographical location (Pérez-Palafox et al., 2022). It is also important to understand that sex ratios in bycatch may not reflect the natural population trends but also fishing selectivity, as fishing gears and nets may disproportionately capture certain size or sex groups (White et al., 2006). Though not significant, the relative size of females of *N. timlei* is larger, hence they may disproportionately get captured.

Although in low numbers, the gravid females were found throughout the years. Four each in the post monsoon and summer season, while 3 and 2 in the pre monsoon and monsoon season, respectively. This shows the absence of a strong seasonal reproductive peak. It aligns with reproductive patterns in electric rays, which exhibit an annual reproductive cycle and prolonged gestation (Devadoss, 1998; Consalvo et al., 2007; El Kamel-Moutalibi et al., 2013). The limited sample size of gravid females (n=13) in the present study restricts us from making certain conclusions. Hence, further sampling would be needed to learn better the reproductive patterns of *N. timlei*.

In our study, the highest number of specimens was observed in post monsoon (33.5%), followed by pre monsoon (25%) and monsoon (23%). The least specimens were recorded in the summer season (18%) (Figure 3).

**Figure 3:**
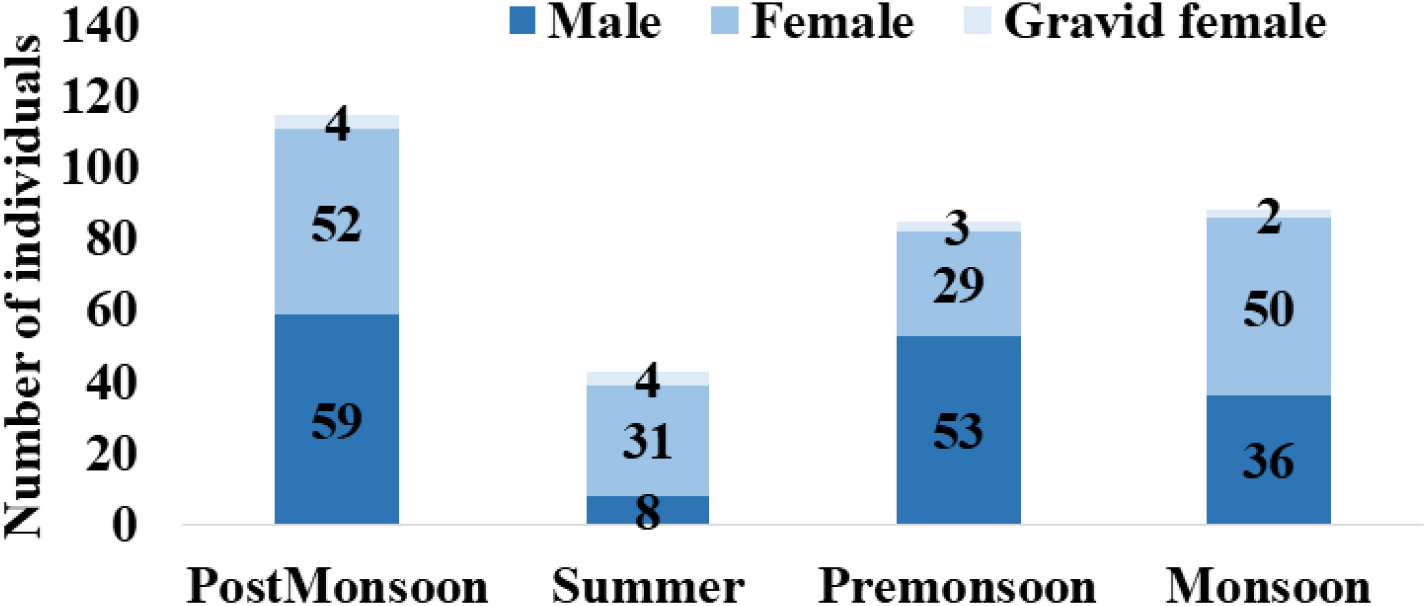
Seasonal frequency in the number of males and females (including gravid females) of *Narcine timlei*.

Though varying frequencies, consistent presence of individuals across seasons, indicate the year-round vulnerability of *N. timlei* to fishing gears. This is concerning for the species, as it is considered a K-selected species due to slow population recovery rates, longer gestation time, and fewer offspring (Dulvy and Forrest 2010). Therefore, bycatch across all seasons can have a significant cumulative impact on population stability.

We observed a variable sex ratio in different seasons: male: female ratio was 1: 0.9, 1: 4.3, 1: 0.60, 1: 1.4, respectively. The summer season had very high female catches, possibly due to females’ reproductive migrations. Similar observations have been reported for *N. brasiliensis* (Rolim, Rotundo and Vaske-Júnior 2016).

### 3.3 Length-Weight Relationship and Growth Patterns

We observed mean weight (mean ± SD) for females as 91.03±39.84, pregnant females as 88.61 ±27.37, and males as 77.82 ±23.49. The mean total length (mean ± SD) was: females 18.21±2.35, pregnant females 18.58 ±1.45, and males 17.54 ±1.66. There was no significant difference observed for the length and weight between males and females. Overall, body size ranged from 14 cm to 29 cm. A large number of specimens, including both females and males, came under the size class of 17-20 cm (57.5%), followed by 14-17 cm (30%). 10% of the specimens were in size class 20-23 cm, and only a fraction of them were larger than 23 cm, mostly females (Figure 4).

**Figure 4:**
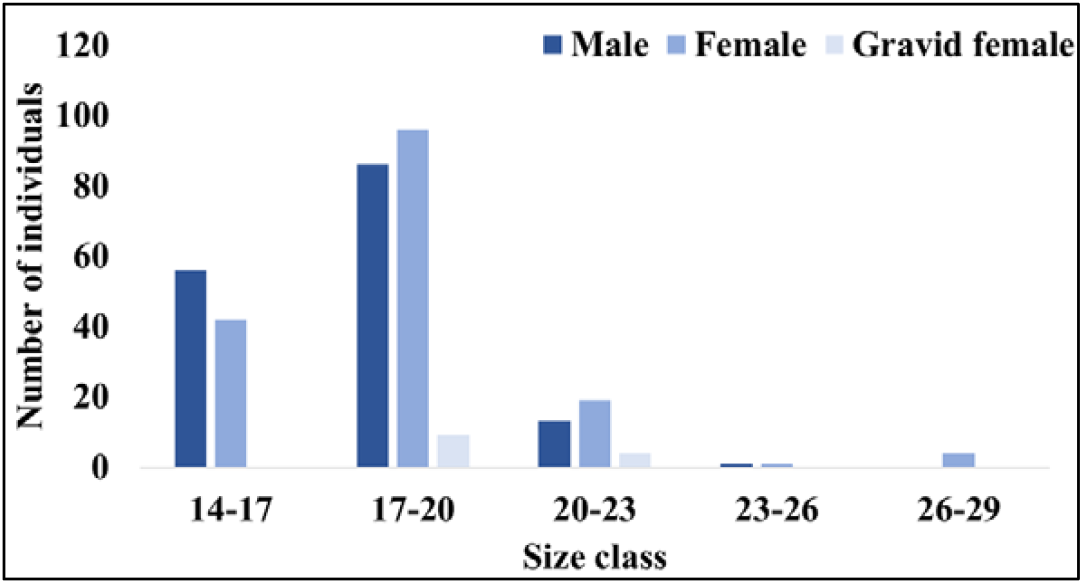
Size distribution of male and female (including gravid females) of *Narcine timlei* The observed average length for *N. timlei* was more than their reported average maturity length of ∼14 cm (Last et al., 2016).

The Length-Weight relationship analysis yielded a b-value of 2.78 (Table 2) for pooled male and female specimens with similar values for females (2.76) and males (2.75) (Figure 5).

**Table 2.** Statistical description and length-weight relationship of *Narcine timlei* collected from the Covelong fish landing centre, Chennai, during the study period.

| Taxa | n | TL (cm) |  | TW (g) |  | regression parameter |  |  |
| --- | --- | --- | --- | --- | --- | --- | --- | --- |
|  |  | Min | Max | Min | Max | a(95%C) | b (95%CL) | r<sup>2</sup> |
| Torpediniformes<br>> Narcinidae ><br>*Narcine timlei*<br>(Bloch &<br>Schneider, 1801) | 33<br>1 | 14.2 | 28.7<br>9 | 38 | 292 | 0.027<br>(0.018-0.038) | 2.78 (2.65-2.90) | 0.84 |

**Figure 5:**
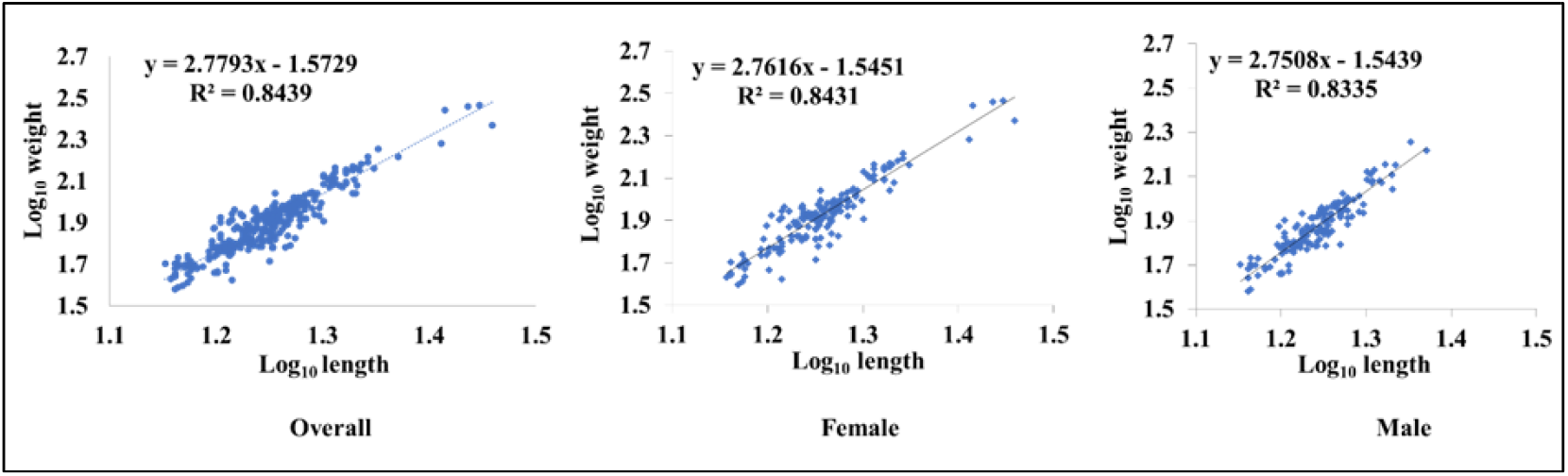
Linear regression analysis of log-transformed length and weight of a) overall, b) Female, and c) Male *Narcine timlei*.

The value suggests allometric growth, where length increases faster than weight. This is a common growth pattern in the other *Narcine* spp. (https://www.fishbase.org/summary/Narcine_brasiliensis.html). The b-value falls within the acceptable limit of 2.5-3.5 for rays (Froese, 2006). The present study contributes significant knowledge on the LWR of *N. timlei*, and r^2^ can be considered near accurate, indicating good predictive power.

### 3.4 Reproductive Biology and Embryo Development

Thirteen gravid females carried a total of 69 embryos (Table 3). Fecundity ranged from 2 to 9 embryos, which is more than the earlier report of 2-3 embryos in each gestation (Last et al., 2016). However, the number remained within the range of other electric rays, such as *Narcine brasiliensis* (Rolim, Rotundo, and Vaske-Júnior 2016), *Torpedo sinuspersici* (Shrikanya and Sujatha, 2014).

**Table 3:**
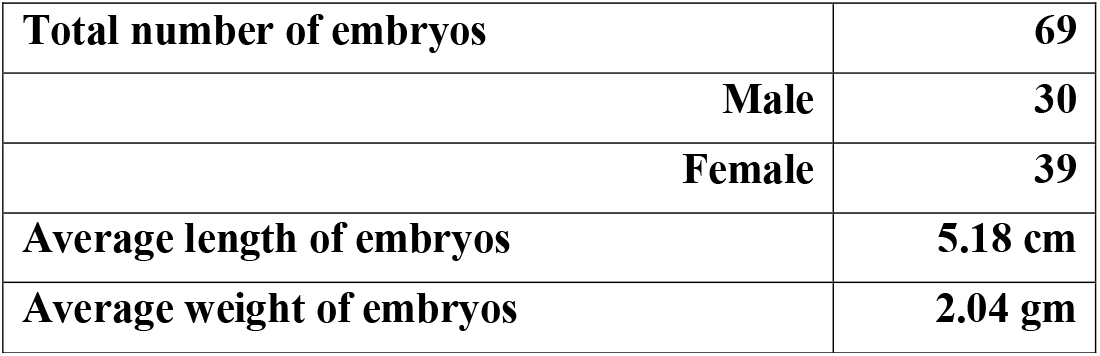
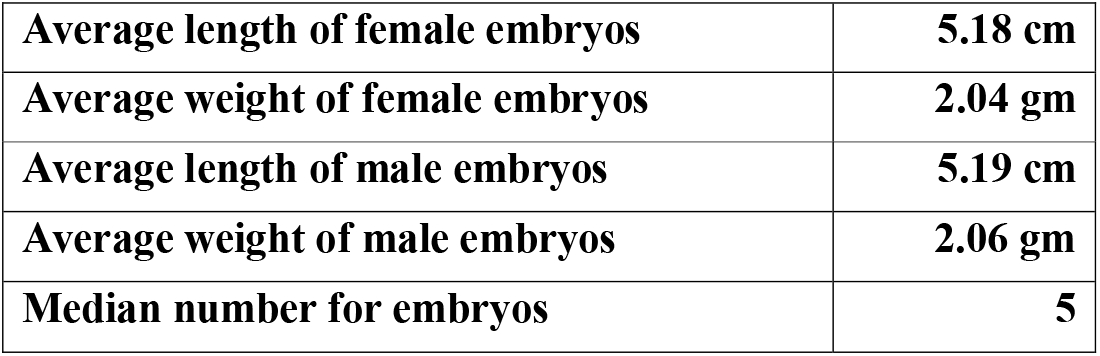
Details of the embryos of *Narcine timlei*.

The observed embryo size was 5.18 ±0.22 cm, and the average weight was 2.02 ± 0.36 g, suggesting mid to almost mature-gestation, as fully developed neonates are almost 6 cm (https://fishbase.se/summary/Narcine-timlei/). Even though we found only 13 pregnant females, we performed the regression analysis for the total length of the mother and number of embryos, but did not find any correlation (R^2^ =0.0036, Figure 6).

**Figure 6:**
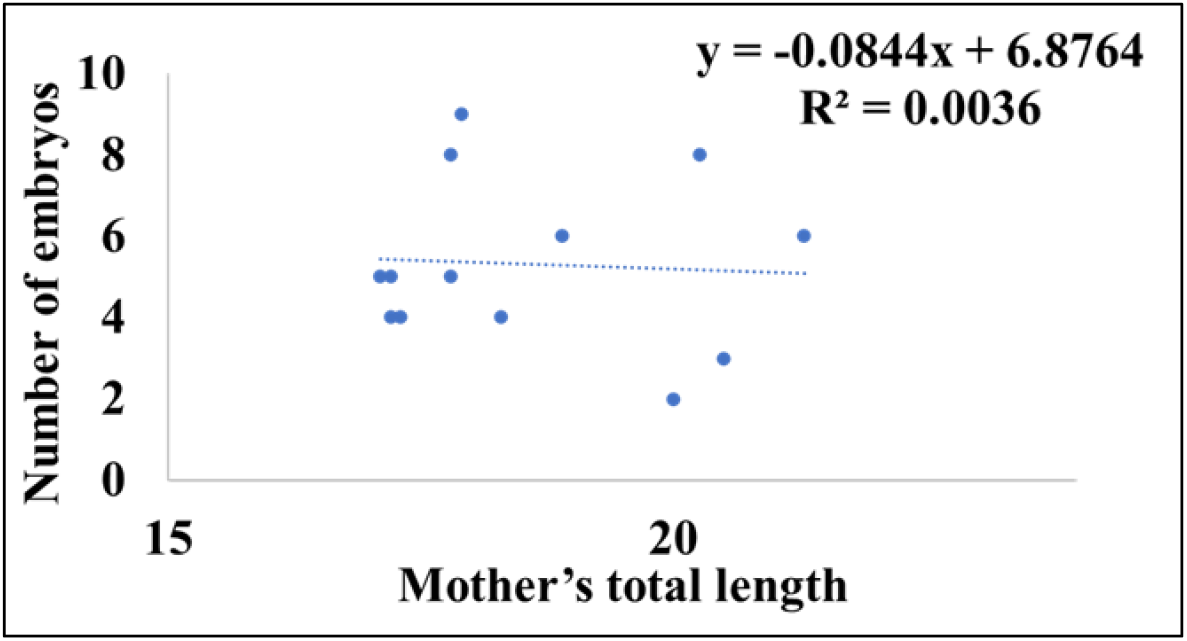
Regression analysis for the total length of the mothers and the number of embryos.

Rolim, Rotundo, and Vaske-Júnior, 2016) also did not find an increase in the litter size with an increase in the total length of the gravid females. However, it is often described that for elasmobranchs, the number of embryos increase with female total length (Walker, 2005). Further sampling is required to ascertain this claim that fecundity does not increase with size in this species.

The length-weight relationships of embryos showed a low r^2^ (0.219) and a b-value of 2.08 (Table 4).

**Table 4.** Length Weight Relationship for the embryos to understand their growth pattern.

| Taxa |  | TL (cm) |  | TW (g) |  | regression parameter |  |  |
| --- | --- | --- | --- | --- | --- | --- | --- | --- |
|  | n | Min | Max | Min | Max | a(95%CL) | b<br>(95%CL) | r<sup>2</sup> |
| *Narcine*<br>*timlei* | 69 | 4.6 | 5.8 | 1.26 | 3.00 | 0.06 (0.014<br>-.30) | 2.08<br>(1.15 -3.00) | 0.21<br>9 |

This indicates a poor predictive fit and strong deviation from the regular growth pattern of *N. timlei*. This may be a regular development pattern during gestation, where embryos prioritise elongation and organogenesis over mass accumulation (Tokunaga et al., 2022). It is important to understand that early development does not conform to conventional growth models and may underline complex intrauterine dynamics.

Interestingly, among the embryos, the skewed sex ratio (1:1.3) in favour of females (30 males and 39 females) was observed. This is almost similar to the sex ratio of adults observed during the study duration (1:1.21). A similar skewed sex ratio (male: female ratio of 1:1.3) was observed in other elasmobranchs, giant electric ray *Narcine entemedor* (Burgos-Vázquez et al., 2017), and guitar fish *Rhinobatus granulatus* (Devodass et al., 1998). Skewed embryo sex ratio could result from maternal physiological regulation or environmental influences. Male and female embryos were found in all seasons; however, the majority of the male embryos (80%) were observed in post-monsoon and summer seasons, while 70% female embryos were observed in post-monsoon and monsoon seasons (Figure 7). This seasonal distribution suggests environmental sex biasing; however, more detailed studies are required to verify the trends.

**Figure 7:**
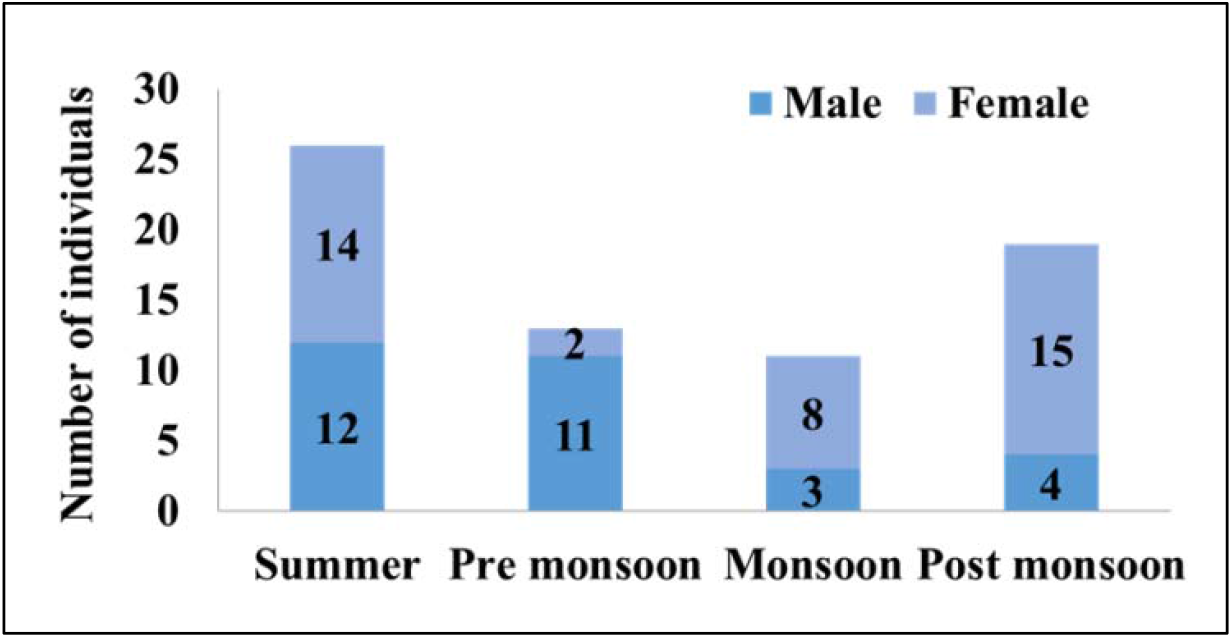
Seasonal frequency in the number of male and female embryos of *Narcine timlei*.

### 3.5 Electric Organ and Electrocyte Morphology

A total of 78 adults and 55 embryos were studied for the morphology of electric organs and electrocytes (Figure 8).

**Figure 8:**
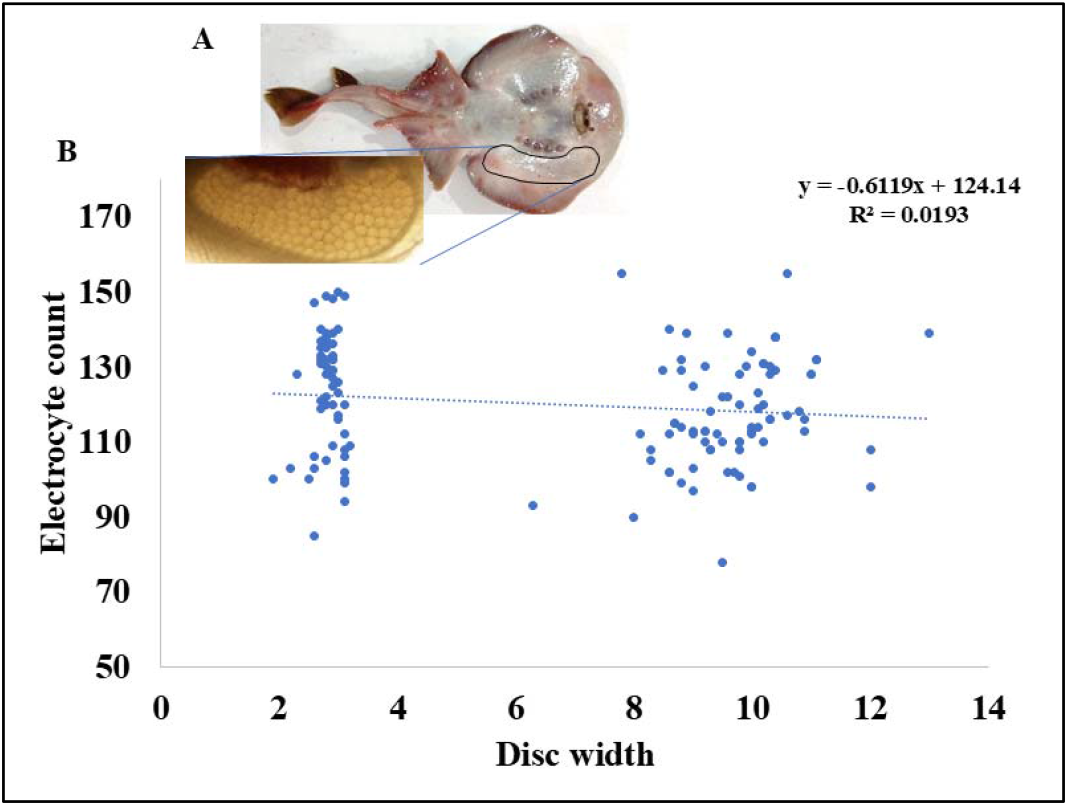
A. Microscopic view of electrocyte cells and electric organ of *Narcine timlei*; B. relationship between Disc-Width and electrocyte count in embryo and adult specimens.

The average length of the electric organ in adults was 4.30 ± 0.47 cm, while in embryos it was 1.22 ± 0.13 cm. The average number of electrolytes in adults was 117.55 ± 14.48, while in embryos it was 123.36 ± 15.92. This result indicates that the electrocyte proliferation happens in the early development, while the size of the cells keeps increasing postnatally. It is possible that before birth, electric organs are at least histologically developed so that functional readiness can be achieved immediately after parturition for their defence (Fox and Richardson 1978).

### 3.6 Ecological and Conservation Implications

We have presented for the first time detailed information on *Narcine timlei*, a small-bodied electric ray, frequently recorded in the bycatches of trawl and gillnet fisheries along the Indian coast. The year-round presence in the bycatch, including the gravid females across seasons, indicates the species’ vulnerability to overfishing. These findings stress the urgent need for monitoring, regulation, and adaptive management strategies. The recently conducted IUCN Red List species assessment did not conduct a study directly on the species but made a conclusion based on the related species and over elasmobranch fisheries data (VanderWright et al., 2021), which may under- or overestimate the risk of this particular species. The present study provides crucial biological and ecological data that can contribute to accurate population evaluations and inform global assessments. The population trend of *N. timlei* is currently declining (https://www.iucnredlist.org/species/161445/178201830), but not enough global efforts are undertaken to reverse the situation. Often, the charismatic species receives more attention, and despite facing equal or greater risks, smaller species are neglected.

At the policy level, Indian has made a national policy on the National Action Plan for Sharks (NPOA-Sharks, 2015) with an aim to promote sustainable management of sharks and rays’ resources, but again the focus was mainly given to commercially important species. Several elasmobranch species found protection under the Indian Wildlife (Protection) Act, 1972, but *Narcine timlei* was not included in the schedule; as a result, unregulated catches are happening and going unmonitored.

## Acknowledgement

We would like to thank The Rufford Foundation, London, for a Small Grant (Reference number: 37306-1). We would also like to thank Sathyabama Institute of Science and Technology, Chennai, for providing the necessary facilities to carry out the lab and field studies. We thank Director (Innovation), Dr. T. Sasipraba, for her constant support and encouragement throughout our research.

## Data availability statement

Data will be shared upon request to authors.

